# AtuR and MarR transcriptional repressors that sense fatty or naphthenic acids and regulate degradation or antimicrobial resistance operons in *Pseudomonas sp.* OST1909

**DOI:** 10.1101/2025.09.24.678253

**Authors:** Tyson Bookout, Jaron Dominquez, Stephanie Wallace, Kira Goff, Steve Shideler, Lisa Gieg, Shawn Lewenza

**Affiliations:** Faculty of Science and Technology, Athabasca University, Athabasca, Alberta, Canada; Department of Microbiology, Immunology and Infectious Diseases, University of Calgary, Calgary, Alberta, Canada; Biological Sciences, University of Calgary, Calgary, Alberta, Canada

## Abstract

Naphthenic acids (NA) are a complex family of acyclic and cyclic carboxylic acids that accumulate in tailings after bitumen extraction from oil sands in Alberta. In response to exposure to naphthenic acids, *Pseudomonas sp*. OST1909 induces the expression of operons regulated by TetR and MarR-type transcriptional repressors. This class of repressor releases from promoters after binding small molecules, leading to a de-repression of the target operons. We propose that these regulators sense fatty and naphthenic acids and control the expression of genes required to cope with their exposure. Plasmid-encoded transcriptional *lux* reporters to the *atu* degradation and *mar* antimicrobial resistance promoters were constructed that also included the corresponding repressor protein, controlled by various strength constitutive promoters. We provide evidence that the *atuAR* and *marR* promoters are directly repressed by AtuR and MarR, respectively. His-tagged MarR and AtuR proteins bound to their promoters containing palindromic binding sites in electrophoretic mobility shift assays (EMSA). Fatty and naphthenic acid compounds and mixtures prevented promoter DNA binding to the repressor proteins in EMSA assays, which suggested direct analyte binding to the protein. The marR-H biosensor construct was functional when introduced into *E. coli*, confirming the role of NA sensing and transcriptional repressor proteins. This study provides mechanistic insight into the use of NA-inducible promoters as bacterial biosensors for NA detection and identifies novel ligands for the MarR and TetR types of repressor proteins.

**Importance:** Understanding the gene expression responses to environmental pollutants is necessary to construct whole cell biosensors that can be employed as a monitoring technology. We have previously identified bacterial genes that were induced by exposure to naphthenic acids, a complex family of cyclic and polycyclic, alkyl substituted carboxylic acids that originate in oil sands but accumulate in the water used to extract bitumen from oil sands. In this study we focus on MarR and TetR-type regulatory proteins and their role in binding fatty acids and naphthenic acids, to further our understanding of how bacteria sense these compounds and mitigate a gene expression response.

## Introduction

TetR and MarR families of transcriptional repressors contribute to the mechanisms by which bacteria sense and respond to environmental changes. TetR family repressors (TFRs) are typically dimeric DNA-binding proteins characterized by a conserved helix-turn-helix (HTH) motif. They function predominantly as repressors by binding to operator sequences within the promoter regions of target genes, thereby blocking RNA polymerase access and preventing transcription. The prototype TetR and MarR repressors regulate antibiotic resistance mechanisms, and diverse processes such as antibiotic biosynthesis genes and nutrient degradation pathways (1). TetR proteins undergo ligand-induced conformational changes that reduces its DNA binding affinity, leading to gene derepression of target genes. There are a diverse number of small molecules sensed by TetR regulators that includes tetracycline, other antibiotics, signalling molecules (acyl homoserine lactones), and compounds involved in amino acid, nitrogen and carbon metabolism (1). The original TetR protein was shown to regulate the *tetA* efflux pump, thereby allowing the cell to detect an antibiotic and induce resistance to that antimicrobial (1).

The MarR family repressors (MFRs) also function as dimeric DNA-binding proteins but exhibit a winged HTH motif (2). Like TFRs, they typically repress transcription by binding to promoter regions and MarR proteins often regulate genes involved in oxidative stress responses, virulence, and antimicrobial resistance. MarR proteins have been shown to sense small molecules such as such as antibiotics, salicylate, reactive oxygen species or phenolic compounds (2, 3). The prototype *Escherichia coli* MarR protein regulates the *marAB* antibiotic resistance operon.

Both MarR and TetR repressors have also been shown to bind and sense fatty acids. TetR homologs such as *Xanthomonas citri* TfmR (4), *Thermus thermophilus* FadR (5), *Bacillus subtilis* FadR (6) and *Pseudomonas aeruginosa* PsrA (7) all bind to fatty acids, leading to the derepression of beta-oxidation and fatty acid degradation pathways. The *Staphylococcus aureus* FarR regulator controls the expression of a resistance-nodulation-division (RND) family efflux pump for fatty acids that is derepressed in the presence of unsaturated fatty acids. Among MFRs, the MarR homolog from psychrophilic bacterium *Paenisporosarcina* sp was purified and shown to bind to palmitic acid, but its overall regulatory function is not yet known (8).

Naphthenic acids (NAs) are a complex mixture of monocyclic, polycyclic, and acyclic alkyl-substituted carboxylic acids typically found in oil sands bitumen, and that accumulate in oil sands process-affected water (OSPW) during oil sands surface mining. Naphthenic acids are the primary compounds of concern that accumulate in tailings, and that ultimately require environmental monitoring and treatment (9, 10). In a previous study, we examined the transcriptome of bacterial genes regulated by the addition of a commercial naphthenic acid mixture, an extract of naphthenic acid fraction compounds (NAFCs) isolated from OSPW, or a simple 9 compound mix of naphthenic and fatty acids (11). Using a *Pseudomonas* isolate recovered from oil sands tailings facilities (12), we identified the genes induced by naphthenic acids exposure. Promoters from naphthenic acid-induced genes were constructed as transcriptional *lux* fusions for the purpose of developing NA biosensors as an environmental monitoring technology (11). The NA-inducible *lux* promoter constructs were confirmed to respond to various model compounds and NA mixtures, where each promoter responded to a unique profile of naphthenic acids. An interesting result of the transcriptome was the induction of numerous metabolite-sensing *tetR* and *marR* regulatory genes, and their adjacent operons in response to naphthenic acids. In this study, we aim to confirm the role of TetR and MarR as repressors and as naphthenic acid sensing proteins.

## Materials and Methods

### Construction of third-generation bioluminescent biosensors

The BioXP 3200™ (CODEX DNA) was used for synthesis of biosensor constructs, which included 1) an NA-inducible promoter to drive expression of the *luxCDABE* reporter in the pMS402 (13) reporter plasmid, 2) a transcriptional repressor gene (*atuR* or *marR*) under the control of various constitutive promoters, and 3) transcriptional terminators downstream of the promoters to prevent read-through (Fig 1). The constitutive promoter controlling the regulator gene expression was either a low, medium or high strength promoter (Table S1, Fig 1). DNA synthesis produced synthetic DNA alone and transformation-ready ligated products into pMS402. For cloning, pMS402 plasmid was linearized by BamHI digestion, and the promoter sequences are synthesized with an extra 30-40 bp on both 5’ and 3’ ends that are complimentary to the sequences flanking the BamHI site. Linear vector and synthetic DNA were mixed with the Gibson assembly mix containing a 5’ exonuclease, DNA polymerase, and ligase. All plasmids were introduced into *Pseudomonas* OST1909 using electroporation.

**Figure 1.**
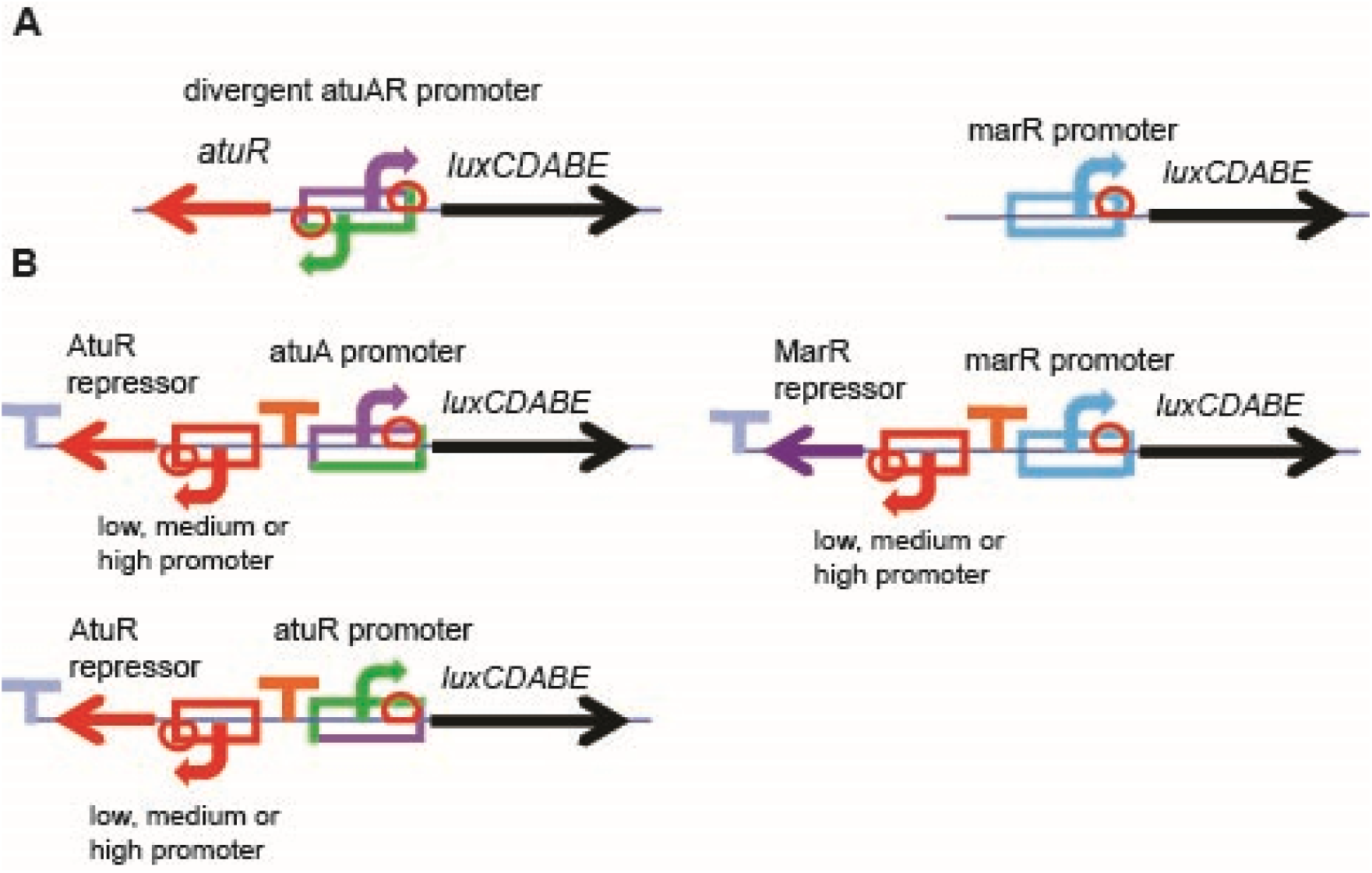
Genetic maps displaying the design of the 3rd generation biosensor constructs. **A)** Maps of the native promoter structure of the *atu* intergenic region containing the divergent *atuA* and *atuR* promoters, and the *marR* promoter in first and second generation, pMS402 plasmid-encoded biosensors. **B)** Third generation, pMS402 plasmid-encoded constructs of the *atuA-lux*, *atuR-lux* and *marR-lux* reporter constructs, which utilize either the “Low”, “Medium”, or “High” strength promoter to control the expression of the AtuR or MarR transcriptional repressor protein. The low strength promoter was upstream of a predicted BCCT family transporter, IH404_RS26360. The medium strength promoter was upstream of a 5-carboxyamino imidazole ribonucleotide mutase, IH404_RS00460. The high strength promoter was upstream of a lipid IV(A) 4-amino-4-deoxy-L-arabinosyltransferase, IH404_RS18330. Terminators (light blue, orange) prevent up or downstream transcription from promoters.

### High-throughput gene expression screening of biosensor constructs for specificity and sensitivity to naphthenic acids

Overnight cultures of biosensors were grown in tubes at room temperature (∼25℃) overnight in 3 mL LB or Terrific Broth (TB) media + 50 µg/mL kanamycin, and subcultured into M9 (Difco^TM^, 248510) or Basal Minimal Medium (BM2) (0.5 mM Mg^2+^) with 20 mM succinate (14) or LB diluted 1/32. The *lux* gene expression assays were conducted in black 96-well, clear bottom plates (Thermo Scientific). NA were diluted to their final concentrations in 99 µl growth media containing 2% DMSO to increase NA solubility and inoculated with 1 µl of the overnight biosensor culture. NA stock solutions were prepared as previously described (11). The three distinct naphthenic acid mixtures were a custom mix of 9 individual model compounds (Table 1; 7 NA compounds and 2 fatty acids), a commercial mix of naphthenic acids containing alkylated cyclopentane carboxylic acids (Sigma-Aldrich, 70340) and a large proportion of acyclic fatty acid compounds (C5-C25) (15, 16), or NAFCs extracted from oil sands process-affected water (17, 18). Naphthenic acid mixtures or model NA compounds were diluted to their final concentrations in growth media containing 2 % DMSO to increase NA solubility. A Breath-Easy® (Sigma-Aldrich) membrane was used to prevent evaporation during a 15-hour protocol in a PerkinElmer 1420 multilabel counter Victor^3^. The plate reader protocol (2 sec shake; gene expression in counts per second (CPS), growth (OD_600_)) included 45 time points, taken every 20 minutes. The gene expression CPS value was typically normalized by dividing the CPS by the OD_600_ for each read. To compare the bioluminescence response to the various compounds tested, the Fold Gene Expression was also calculated by dividing the CPS/OD_600_ values of the NA treated sample by those of the untreated sample. A fold change of 1 indicates no change, and a fold change of 2 indicates a doubling of *lux* expression in comparison to the untreated control.

**Table 1.** Components of the custom 9x naphthenic acid mixture (10 mg/ml)

| Compound | Abbrev. | Final Concentration |
| --- | --- | --- |
| Hexanoic acid | HA | 1.11 mg/ml |
| Decanoic acid | DA | 1.11 mg/ml |
| Cyclopentane carboxylic acid | CPCA | 1.11 mg/ml |
| Cyclohexane pentanoic acid | CHPA | 1.11 mg/ml |
| Cyclohexane acetic acid | CHAA | 1.11 mg/ml |
| 3-Phenylpropionic acid | PPA | 1.11 mg/ml |
| 4-Dimethylphenylacetic acid | DMPAA | 1.11 mg/ml |
| 1-Adamantane carboxylic acid | ACA | 1.11 mg/ml |
| 5,6,7,8-Tetrahydro-2-naphthoic acid | THNA | 1.11 mg/ml |

### PCR and Gibson Assembly cloning of the His_6_-tagged AtuR and MarR proteins

The *atuR* and *marR* repressor genes were cloned as C-terminal His_6_-tagged proteins into the IPTG-inducible expression vector pMCSG59 (19) using the benchtop Gibson Assembly kit (New England BioLabs, E5510S). The vector contained a TEV cleavage site upstream from the His_6_-tag, for the removal of the histidine residues after purification. OST1909 genomic DNA from was purified and used as the template for PCR amplification of both *atuR* and *marR*. The PCR primers designed for the amplification of the regulator genes are PsAtuR-F/R, and PsMarR-F/R (Table S1). These primers were designed to contain 20 bp matching the gene of interest with the addition of a 5’ 30 bp extension complimentary to vector SmaI site, which allows efficient and unidirectional cloning into the vector through Gibson Assembly. The pMCSG59 vector was linearized with blunt-end SmaI digestion (50 ng) and mixed with 25 ng of either the *marR* or *atuR* amplified insert and 10 µl of the Gibson Assembly mix in a final volume of 20 µl and incubated at 50 ℃ for 30 minutes. Ligation products were transformed into chemically competent *E. coli* BL21 Gold using heat shock and plated on LB + 100 mg/L carbenicillin. Transformed colonies were verified through colony PCR using original primers (Table S1).

### Purification of His-tagged transcription factors

AtuR-His_6_ and MarR-His_6_ were purified using nickel column chromatography Overnight cultures of protein expression strains were sub-cultured in 1 L of LB + 100 µg/ml carbenicillin (4L flask) and incubated at 37 ℃ until they reached a mid-log OD_600_ of 0.6 - 0.8. After cooling to 18℃, 1 mM IPTG was added and flasks were incubated overnight at 18 ℃ with 175 rpm shaking, and cells were harvested by centrifugation. The pellet was suspended in 40 ml Binding buffer, with the addition of 400 µl of 100X protease inhibitor cocktail (PIC) (100 µM benzamidine HCL, and 50 µM phenylmethylsulfonyl fluoride in 95% ethanol), and cells were lysed by sonication. The sonicated sample was clarified and incubated with Ni-NTA beads at 4 ℃ for 60 minutes.

After incubation with Ni-NTA, samples were centrifuged (5 min, 800 rpm, 4 ℃), washed twice with 25 ml Wash buffer and 250 µl of 100X PIC, with a final resuspension in 25 ml Wash buffer without PIC. The resuspended resin was poured into a column, flowthrough was collected, and 10 ml Elution buffer was used to collect the protein of interest. Elution was monitored for the presence of protein using Bradford reagent and SDS-PAGE. To remove any histidine containing proteins from the Ni-NTA matrix, the TEV protease (30 µg) was used to cleave the His_6_-tagged protein in dialysis tubing overnight, then loaded directly onto a new Ni-NTA slurry and run twice through the column twice to remove contaminants. Proteins were quantitated with the NanoDrop, resuspended in 1 mM ETDA and 2 mM DTT and visualized on Coomassie Blue stained SDS-page gels made with a 12% resolving gel and a 5% stacking gel.

### Electrophoretic Mobility Shift Assays

The AtuR and MarR transcription factors were examined for mobility shifts upon binding to their target DNA sequences, where increasing amounts of the purified protein (0.6 to 4.8 µg) were incubated with 100 ng of either the entire PCR amplified promoter region or a specific 30 bp DNA probe identified using the EMBOSS palindrome finder (20). Both strands of each ∼30 bp probe were ordered as single-stranded oligos and annealed by mixing 12.5 µl of both strands and decreasing the temperature from 95 ℃ to 25 ℃ over 45 minutes. DNA probes are shown with underlined regions indicating inverted repeats and the primers used for amplification of the promoter region can be found in Table S1. The repressor proteins were incubated with DNA, in the presence or absence of exogenous naphthenic acid model compounds or mixtures, varying the amount of protein and NA, in a 40 µl of reaction buffer containing 0.02 M Tris-HCl and 5 mM MgCl_2_ for 1 hour at room temperature (21). Protein, DNA, and NA samples were run on 5% polyacrylamide non-denaturing gels and the DNA bands stained by submerging the gel in a SYBR-Safe staining solution and visualized under UV light.

## Results

### Design of transcriptional *lux* reporters containing NA-inducible promoters and their transcriptional repressor proteins

We recently identified naphthenic acid-induced genes with RNA-seq transcriptome experiments and validated these expression patterns using transcriptional *luxCDABE* reporters (11). The transcriptional *lux* fusions encoded on plasmids enabled strains carrying these plasmids to function as biosensors for the detection of model NA compounds, as well as custom and commercial NA mixtures, extracts of NAFCs in bacterial culture conditions and in raw OSPW samples (11). In the first and second generation of biosensor constructs, only the NA-inducible promoter was cloned into the plasmid-encoded *lux* reporter plasmid (11). Here we report the construction and testing of third generation constructs that contain 1) transcriptional *lux* reporters and 2) the predicted transcriptional repressors that are hypothesized to control the expression of these NA-inducible promoters (**Fig 1**).

### The *atuA* and *atuR* promoters respond to acyclic naphthenic acids and are repressed by AtuR

The divergent *atuA* and *atuR* promoters in *Pseudomonas sp*. OST19019 were previously shown to respond to a Sigma-Aldrich commercial NA mixture that contains alkylated cyclopentane carboxylic acids and a large proportion of acyclic fatty acid compounds (C5-C25) (15, 16). The promoters also responded specifically to individual model compounds of saturated and branched fatty acids, including hexanoic acid (HA), pentadecanoic acid (PA), stearic acid (SA) and citronellic acid (CA) (6-18 carbons) but not to NA compounds (11). The OST1909 *atuABCDEF* gene cluster encodes a probable degradation operon that metabolizes saturated and branched fatty acids (**Fig 2**), given its similarity to the *atuABCDEFGH* operon was originally identified in *Pseudomonas aeruginos*a, which is required for growth and degradation of branched fatty acids, or acyclic terpenes (22–24).

**Figure 2.**
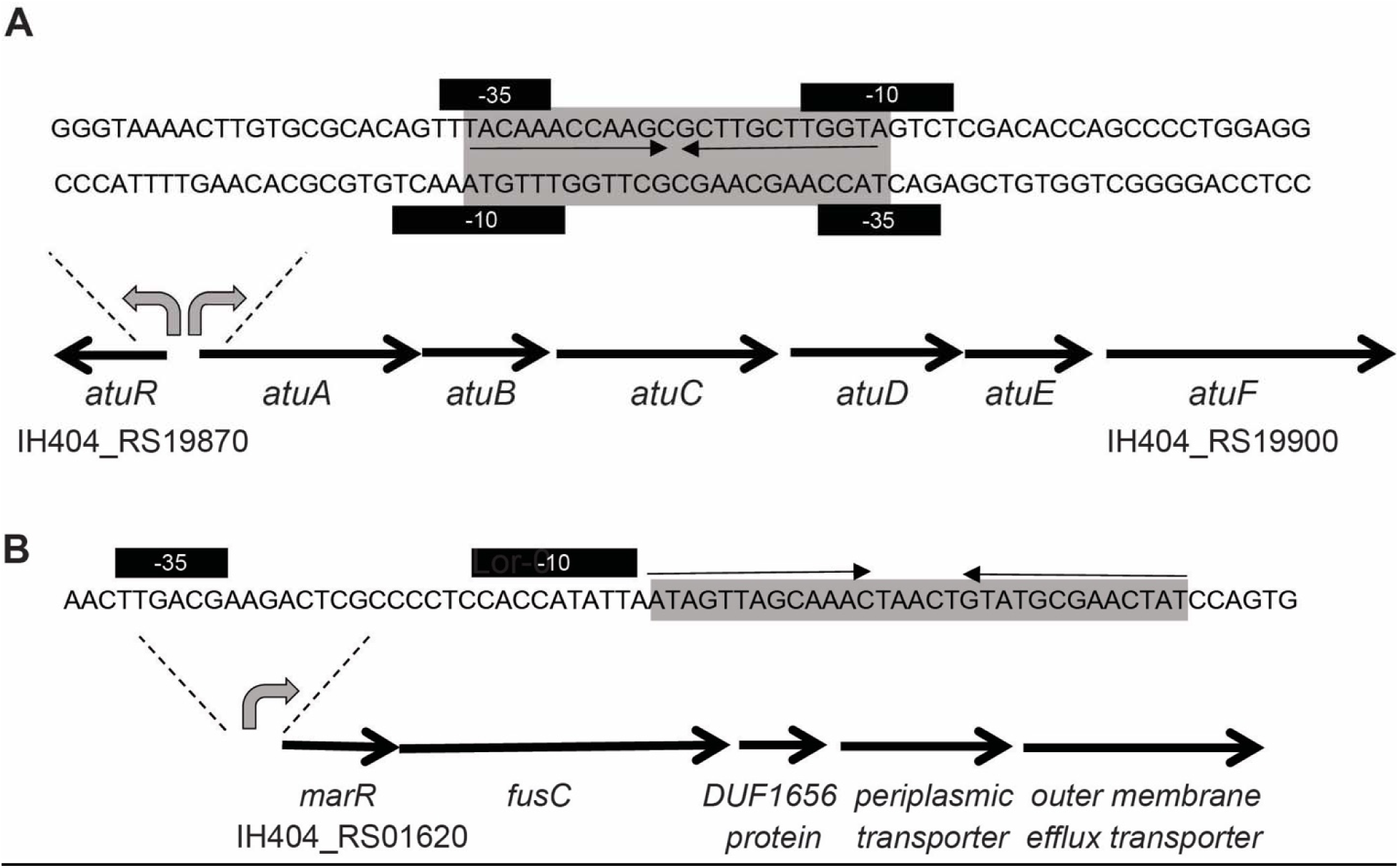
Predicted palindrome sites in promoters regulated by the AtuR and MarR repressors. **A)** The *atu* cluster was originally identified as degradation genes required for acyclic terpene utilization (*atu*). Between the regulator and degradation genes there are divergent promoters containing inverted repeats that overlap the −10 and −35 elements for both *atuA* and *atuR* promoters. **B)** The *marR* regulator gene is within an operon annotated as fusaric acid resistance and efflux genes. This promoter contains inverted repeats upstream of the −10 and −35 elements. Sigma 70 promoters (black boxes) were predicted using BPROM and the operator sites consisting of inverted repeats (grey boxes) suggesting TetR and MarR binding. Operator sites are downstream or overlap the promoter, which prevents RNA transcription.

Here we designed the 3^rd^ generation *atuA-lux* biosensor constructs with various strength promoters to control the expression of the respective transcriptional repressor (Fig 1). The three OST1909 promoters selected to control the repressor gene expression were shown to have either <u>L</u>ow, <u>M</u>edium, or <u>H</u>igh promoter activity (Fig 3A-C). To confirm that these promoters were constitutive, we showed that expression was not changed when *Pseudomonas sp. OST1909* cultures were exposed to a commercial NA mixture containing a large proportion of acyclic compounds (Sigma-Aldrich), or a custom mix of 9 NA compounds (Fig 3A-B). Therefore, these promoters were cloned upstream of the *atuR* or *marR* regulator gene, in order to produce constitutive levels of repressor protein at low, medium and high amounts in the cell. We hypothesized that increasing expression of the repressor protein would lead to decreasing levels of target gene expression.

**Figure 3.**
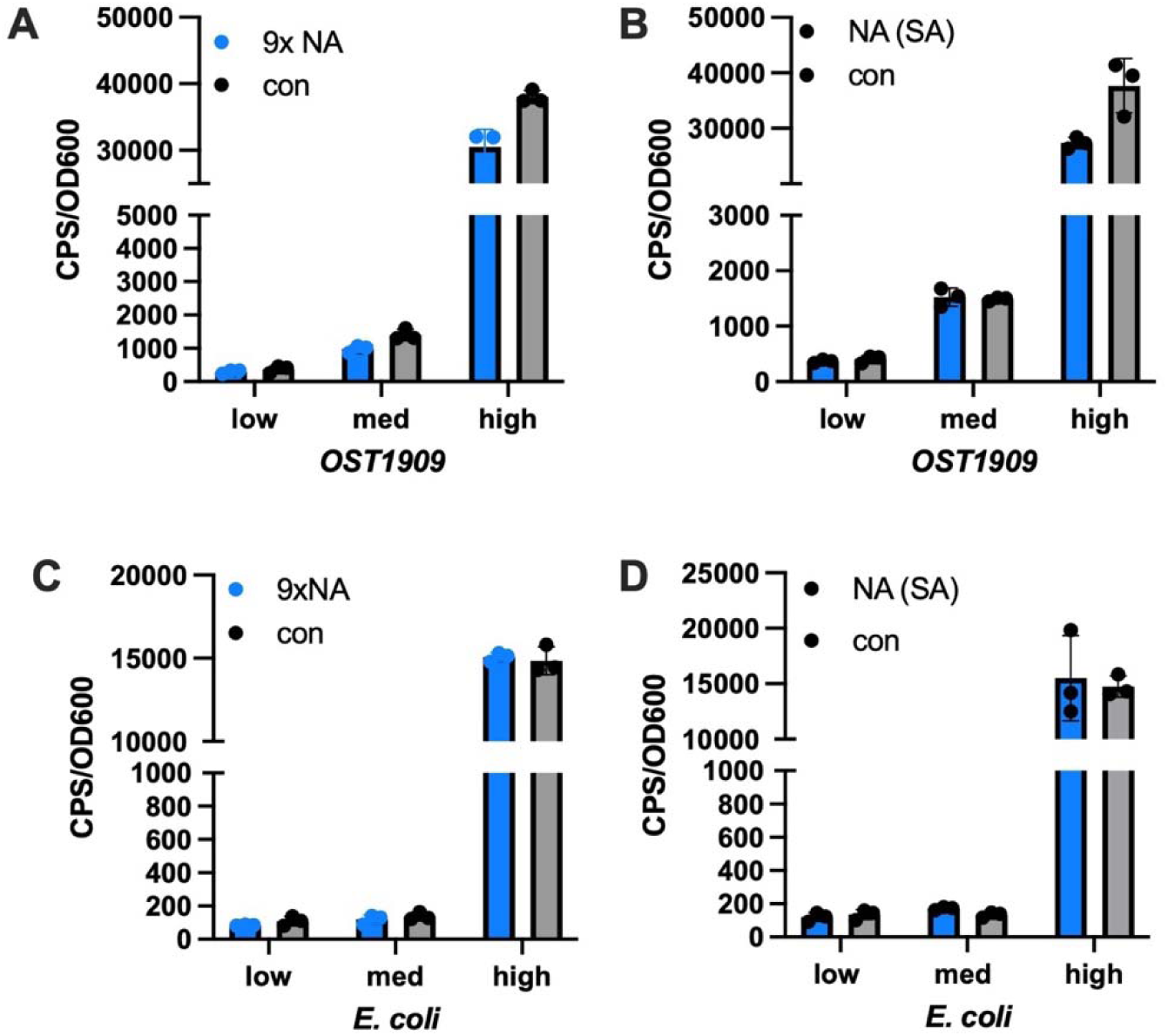
Expression of low, medium and high strength, constitutive promoters. The pMS402 plasmid-encoded transcriptional *lux* fusions were expressed in *Pseudomonas sp.* OST1909 **(A,B)** and *E. coli* DH5α **(C,D)** and maximum gene expression was reported at the late log stage of growth (t = 7rs), in the absence or presence of 100 mg/L commercial NA mixture (Sigma Aldrich, SA) or a custom 9xNA mix. Experiments were performed in 1/32 LB, all values shown are the average of triplicates plus standard deviation, and each experiment was performed 3 times.

In Figure 4, we demonstrated that both the *atuA* and *atuR* promoters were inversely regulated depending on the level of repressor protein expression. Under non-inducing conditions, when the repressor was mostly highly expressed, this resulted in strong repression of basal *atuA* and *atuR* expression. Conversely, when the repressor was expressed from a weak or low strength promoter, the *atuA* and *atuR* promoters had the highest level of basal expression (Fig 4 A,D). These data support the role of AtuR as a transcriptional repressor of both divergent *atuA* and *atuR* promoters within this intergenic region. When calculating the fold induction values of the *atuA-lux* constructs after exposure to a commercial mixture of NA, the biosensor construct with Medium strength expression of AtuR demonstrated the highest fold induction (∼10-fold) and was therefore the most sensitive in responding to the commercial mixture of NA (Fig 4C). In agreement with this gene expression data, the intergenic promoter contains a single predicted AtuR inverted repeat, binding site that is positioned overlapping and in between the predicted −10 and −35 regions of the RNA polymerase binding site of both promoters (Fig 2).

**Figure 4.**
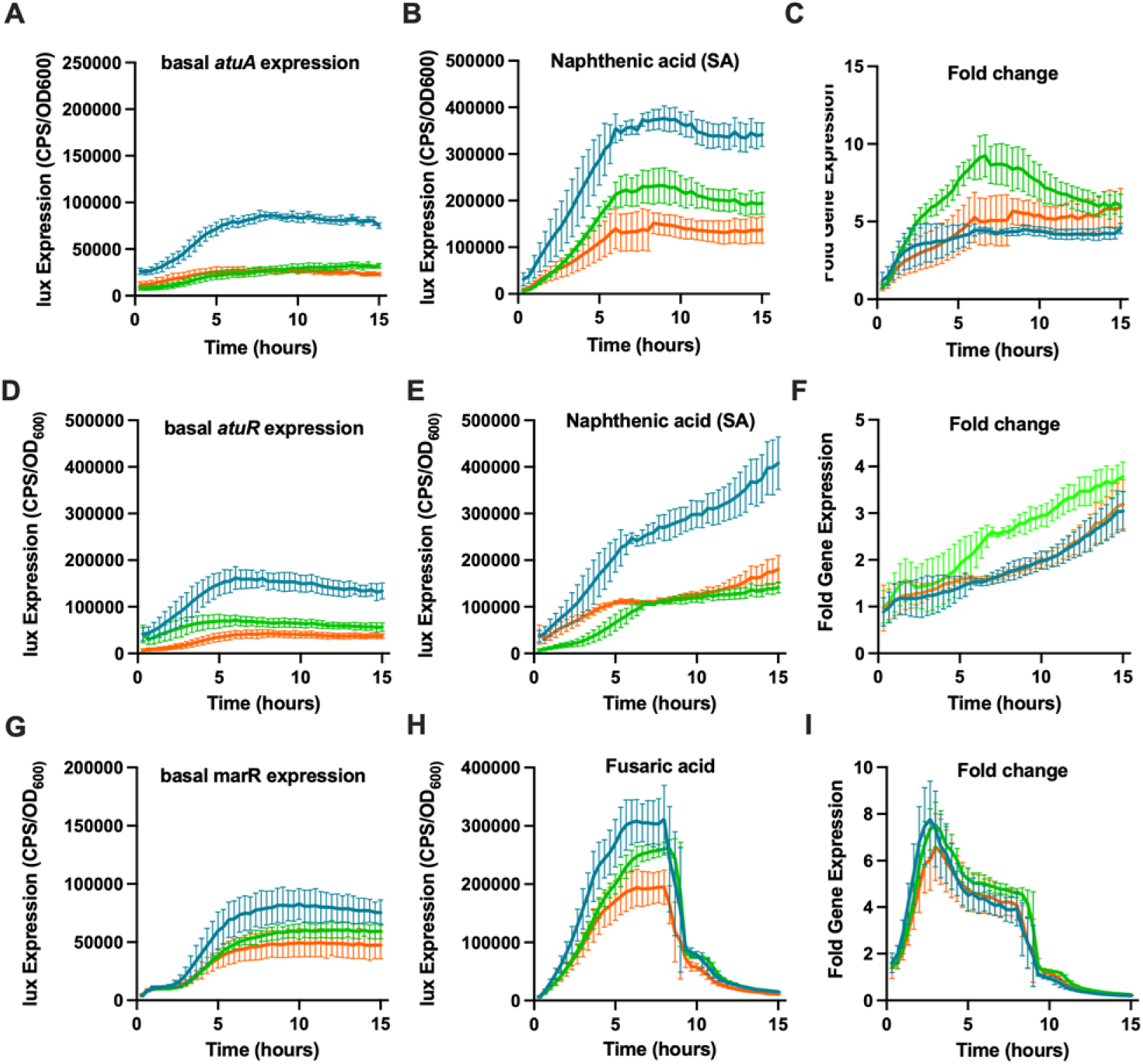
Gene expression from third generation constructs of *atuA-lux*, *atuR-lux* and *marR-lux* transcriptional reporters. **A)** *Basal atuA-lux,* **D)** *atuR-lux* and **G)** *marR-lux* expression under non-inducing conditions. Expression under inducing conditions by exposure to 400 μg/ml commercial naphthenic acid mixtures (Sigma Aldrich, SA) **(B,E)** or 25 μg/ml fusaric acid **(H)**. The fold change induction after NA exposure **(C,F,I).** The three constructs varied due to constitutive expression of the AtuR or MarR repressor from high (orange), medium (green) or low (blue) strength promoters. Gene expression at each time point was expressed as CPS (counts per second) that was normalized by the OD_600_ (growth) at 20 min intervals throughout a 15 hr period, with standard deviation values.

The various *atuR-lux* constructs were also tested for their ability to respond to a commercial mixture of NA. Basal *atuR* expression was highest when the repressor was expressed from the Low strength promoter. Under inducing conditions, the *atuR* promoter was most strongly upregulated in the construct where the AtuR repressor was expressed at medium levels (Fig 4B,E). Ultimately, the biosensors with the highest fold changes were observed when the repressor was under the control of a medium strength promoter (Fig 4C, F). In conclusion, the *atuA* promoter was the most highly responsive to this commercial NA mixture, was maximally induced at an earlier time point and is the preferred promoter for detecting this NA mixture (Fig 4, Table 2). We also compared expression of the low, medium and high constitutive promoters to the native, inducible promoter that controls the *atuR* repressor. In this comparison, the ‘wild type’ dual *atuA-atuR* promoter had even higher fold changes and therefore sensitivity, than either of the synthetic variants (Table 2). These results demonstrate that varying the expression of the repressor protein affects both basal and induced expression, consistent with the role of AtuR as a repressor.

**Table 2.** Comparison of sensitivity of various *atuA*, *atuR* and *marR* biosensor constructs.

| AtuR level of expression | *atuA-lux* |  | *atuR-lux* |  | *marR-lux* |  |
| --- | --- | --- | --- | --- | --- | --- |
|  | *Max* | Time | *Max* | Time | *Max* | Time |
|  | Induction<sup>a</sup> | (hrs) | Induction<sup>a</sup> | (hrs) | Induction<sup>b</sup> | (hrs) |
| Constitutive Low | 4.5 | 6.6 | 3 | 15 | **7.8** | 2.7 |
| Constitutive Medium | **9.2** | 6.6 | 3.7 | 15 | 7.5 | 3 |
| Constitutive High | 5.5 | 7.3 | 3.2 | 15 | 6.6 | 3 |
| Inducible Wild type | 13 | 5 | **4** | 15 | N/A | N/A |

<sup>a</sup> *atuA* and *atuR* constructs response to 400 µg /ml commercial naphthenic acids (SA).
<sup>b</sup> *marR* constructs response to 25 µg /ml fusaric acid.

### The *marR* promoter responds to fusaric acid and is repressed by MarR

Fusaric acid (FA) is a fungal produced antimicrobial that has a naphthenic acid structure with a nitrogen-containing ring structure (25). The *marR* operon encodes genes annotated as fusaric acid resistance genes, including *fusC* and components of an RND efflux operon (Fig 2). FA resistance increases when FusC and nearby genes are expressed or induced in other bacteria (26, 27). The *marR* promoter responds to fusaric acid, but also naphthenic acid model compounds with classic or complex structures, raw OSPW samples and to NAFCs extracted from oil sands tailings water (11). When comparing the various synthetic *marR* biosensor constructs response to fusaric acid (Fig 4D-F), the *marR* promoter demonstrated the highest expression level when the MarR regulator was expressed from a low constitutive promoter. As the strength of promoter increased that controls the MarR repressor, there was a decreased expression from the *marR-lux* reporter (Figure 4D-F), which was similar to the pattern of the AtuR-repressed promoters (Figure 4A-F).

Under inducing conditions in the presence of 25 μg/ml of fusaric acid, the highest level of expression was again seen when the repressor was expressed at low levels (Fig 4H). In general, higher levels of repressor expression resulted in lower levels of target gene expression, which supports the hypothesis that MarR is a repressor. When the fold induction values were determined, the low and medium level repressor constructs showed the highest inducibility, and therefore sensitivity to fusaric acid.

### Bioinformatic prediction and validation of a novel AtuR-regulated, fatty acid degradation gene

We searched the OST1909 genome using BLASTN analysis with the 24 bp, imperfect inverted repeat AtuR binding site in the *atuAR* divergent promoter (5’ -*TACAAACCAAGC*GCTTGCTTGGTA) and identified one other promoter containing a possible repressor site from the gene IH404_RS20745 that encodes an acyl-CoA dehydrogenase family protein (18/24 matches), which are enzymes that participate in beta-oxidation of fatty acids. The inverted repeat site overlaps a predicted sigma 70 promoter in this case (Fig 5). Due to the presence of an AtuR repressor site in the promoter of a gene that appears to be involved in fatty acid degradation (Fig 2), we cloned this promoter as a transcriptional *lux* reporter to test if the promoter was inducible by naphthenic acids. Figure 5 demonstrates that the p*20745-lux* reporter is induced by the commercial naphthenic acids mix (Sigma-Aldrich), as well as to individual fatty acids citronellic and decanoic acid, which is the same specificity as the *atuA* promoter (11). This indicates that AtuR controls two fatty acid degradation gene promoters among its regulon. We also searched the OST1909 genome for possible MarR inverted repeat binding sites but there was only one site in the promoter controlling the *marR* regulator (5’-*ATAGTTAGCAAAC*TAACTGTATGCGAACTAT), suggesting one operon in the regulon that encodes the fusaric acid resistance genes.

**Figure 5.**
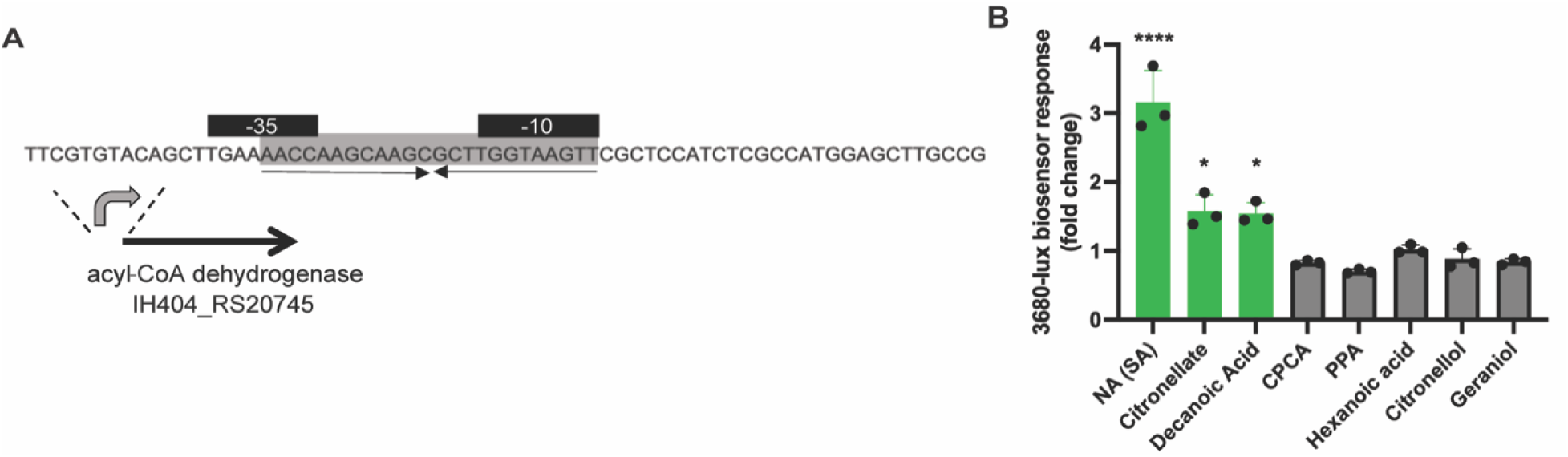
An AtuR binding site is predicted in a putative fatty degradation gene encoding an acyl CoA dehydrogenase. **A)** An acyl-CoA dehydrogenase contains a promoter with a predicted AtuR binding site (grey box, inverted repeats) that overlap the −10 and −35 elements of the IH404_RS20745 promoter. **B)** Gene expression from a p20745-*lux* transcriptional reporter was induced by exposure to 200 mg/L commercial NA mix (Sigma Aldrich), 100 mg/L of citronellic acid and decanoic acid, but not cyclopentane carboxylic acid, 3-phenylpropionic acid, hexanoic acid, citronellol or geraniol. A one-way ANOVA revealed a statistically significant difference between treatment groups (F (8, 18) = 48.63, p<0.0001). A post hoc Dunnett’s test for multiple comparisons was conducted between each compound or mix to test significant relationships between each group (α = 0.05, n = 3). Treatment groups with significant results compared to the control are indicated with asterisks and green bars (**** p<0.0001, * p<0.05). All gene expression values shown represent the average of triplicates with standard deviation and each experiment was performed 3 times.

### Binding of MarR and AtuR to upstream DNA regions containing predicted promoters with inverted repeat operator sites

To confirm the direct binding of these transcriptional repressors to their predicted operator sites, MarR and AtuR were purified and His_6_-tagged proteins and used in electrophoretic mobility shift assays (EMSA). These experiments involved the incubation of either AtuR or MarR with its respective promoter region. The non-denaturing polyacrylamide gels in Figure 6 display the mobility shifts of the fluorescent promoter DNA after incubation with increasing concentrations of its respective repressor protein. We observed a predicted upshift in size of the DNA band with as little as 2.4 ug of repressor protein, which was the result of a protein-DNA complex for both AtuR and MarR (Fig 6). The binding of MarR and AtuR to its promoter DNA was incomplete, and despite extensive troubleshooting efforts, complete binding and upshifting of all stained DNA was not observed.

**Figure 6.**
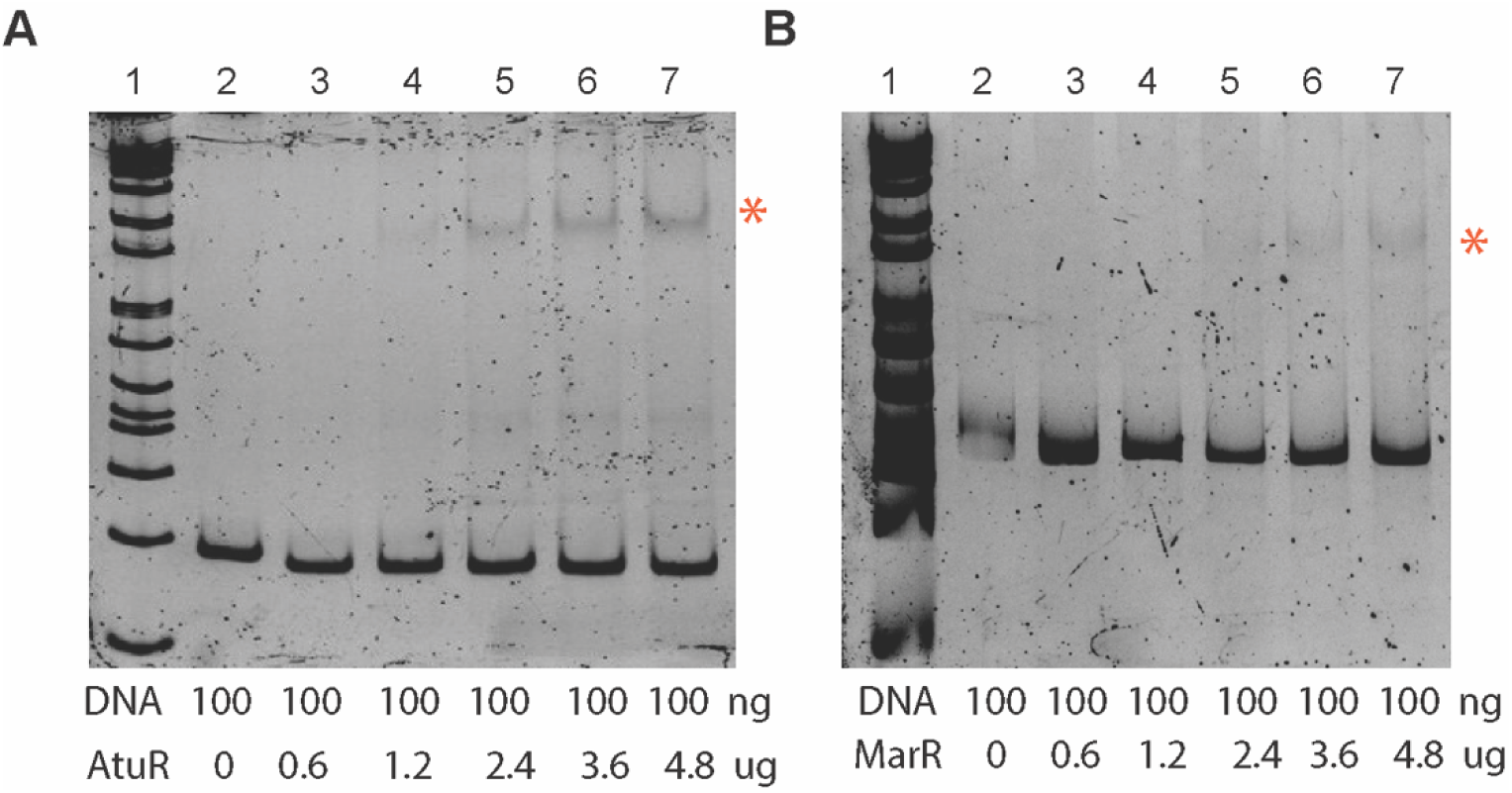
AtuR and MarR binding to their respective promoter DNA. EMSA gel demonstrating the binding of **A)** AtuR-His_6_ to a 151 bp upstream DNA region and **B)** MarR-His_6_ to its 415 bp upstream DNA region. Orange asterisks highlight the shifted size of a regulatory protein-DNA complex. Increasing concentrations of protein (µg) were incubated with a 100 ng of promoter DNA, which was stained for visualization with SYBR, with the 1 kb plus DNA ladder on the left. Each experiment was performed at least 3 times.

### Naphthenic acid and fatty acid compounds prevent binding of AtuR to its promoter DNA

AtuR is one of 27 TetR family regulators in the *Pseudomonas sp*. OST1909 genome. While many TetR homologs can sense fatty acids (7, 5, 6, 28), here we explore the ability of AtuR to bind to both fatty and naphthenic acids. Figure 7A (lanes 1 and 2) depicts binding of AtuR to its promoter and examines the influence of adding naphthenic acids as a third component into the binding mixture. When DNA (100 ng) and AtuR (4.8 µg) are incubated alone, there are two new bands with an increased mass, which may represent the binding of monomeric and dimeric AtuR to its promoter (Fig 7A, lane 2). The binding of AtuR to its promoter DNA was inhibited by decanoic acid (DA), model NA compounds including cyclopentane carboxylic acid (CPCA), cyclohexane pentanoic acid (CHPA), 3-phenylpropionic acid (PPA), adamantane carboxylic acid (ACA), fusaric acid (FA), and partially inhibited by a commercial NA mix (Fig 7A).

**Figure 7.**
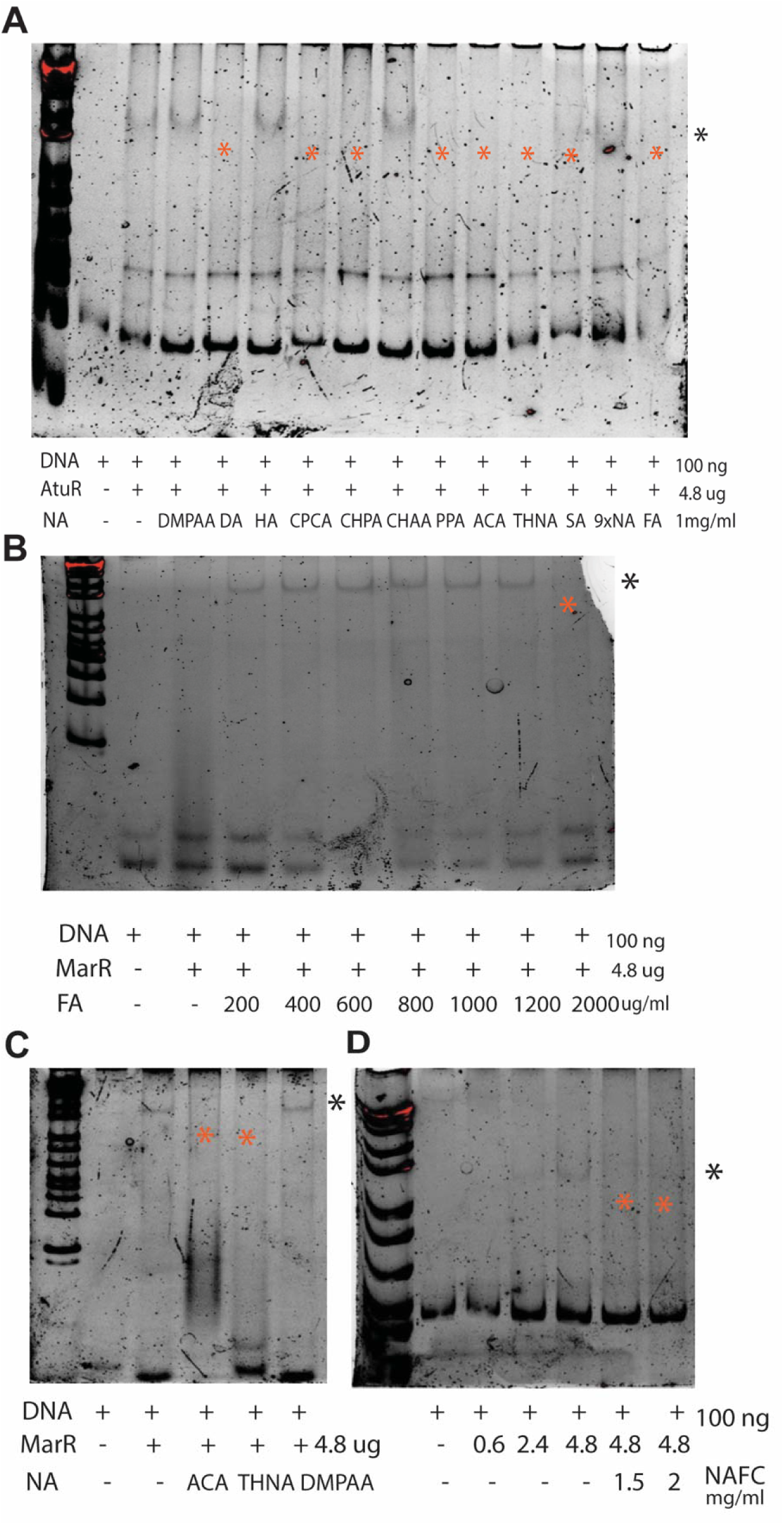
Naphthenic acids prevent AtuR and MarR binding to promoter DNA. **A)** EMSA gel demonstrating the binding of 4.8 µg AtuR to 100 ng promoter region, as well as to 1 mg/ml (0.1%) of different NA compounds. The following naphthenic acid compounds were added at 1mg/ml: 4-dimethyl phenylacetic acid (DMPAA), decanoic acid (DA), hexanoic acid (HA), cyclopentane carboxylic acid (CPCA), cyclohexane pentanoic acid (CHPA), cyclohexane acetic acid (CHAA), 3-phenylpropionic acid (PPA), adamantane carboxylic acid (ACA), 5,6,7,8-tetrahydroic 2-naphthoic acid (THNA), commercial NA mixture from Sigma Aldrich (SA), 9xNA custom mixture (9xNA) and fusaric acid (FA). **B)** EMSA gel demonstrating the binding of 4.8 µg MarR to 100 ng of the 30 bp inverted repeat region of the *marR* promoter, and with the addition of increasing amounts of fusaric acid (FA). **C**) MarR binding to the 30 bp inverted repeat region of the marR promoter, with the addition of ACA, THNA or DMPAA. **D)** MarR binding to its promoter DNA region (415 bp), with the addition of naphthenic acid fraction compounds (NAFCs). The presence and concentration of DNA, MarR and NA are indicted in the legend beneath each figure. The black asterisks indicate the locations for the migration of the AtuR-DNA complexes, and the orange asterisks indicate the absence of the upper size shifted AtuR-DNA complex. Each experiment was performed at least 3 times.

This broad naphthenic acid binding ability was unexpected, as we predicted that the strongest NA binding would occur with acyclic fatty acid compounds, as this promoter is induced mainly by acyclic compounds (11). Binding of decanoic acid and the commercial NA mix was consistent with the biosensor responses, which is an indicator of what compounds bound to AtuR in the bacterial cytoplasm (Fig 7A). This in vitro binding assay provides evidence that a broad spectrum of NA compounds can prevent AtuR binding to DNA, indicating AtuR undergoes a confirmation change after NA binding and release from its promoter. Binding of model NA compounds in the EMSA may be due to high concentrations of NA used (1 mg/ml), a level which is likely not achieved in the cytoplasm, or in tailings water.

### MarR binds to complex naphthenic acids compounds and those extracted from OSPW

The protein IH404_RS01620 is one of eight MarR family regulators present in *Pseudomonas sp*. OST1909. MarR type proteins function in a comparable manner to TetR proteins and have been reported to regulate processes that include antibiotic resistance, among others (3). The *marR* containing operon in OST1909 (Fig 2) is highly induced by naphthenic acid compounds with classic and complex structures and also induced by the naphthenic acid fractions compounds (NAFCs) that are extracted from oil sands tailings facilities (11). Given the ability to MarR proteins to recognize small molecules, we predicted that naphthenic acid compounds that induce expression of the *marR* operon are detected inside the cell, leading to a loss of promoter binding, and induction of the operon.

In Figure 7B, the binding of MarR to a short, double oligonucleotide sequence that contains the predicted binding site is demonstrated. A gradient of fusaric acid concentrations was added to attempt to prevent MarR binding of DNA, and only the highest concentration of fusaric acids (2000 mg/ml) was capable of preventing MarR from binding to its repressor site (Fig 7B), which is consistent with fusaric acid acting as an inducer of this operon (11). In Figure 7C, MarR binding to its operator DNA site was inhibited by in the presence of adamantane carboxylic acid (ACA) or 5,6,7,8-tetrahydro-2-napthoic acid (THNA), both of which are also compounds that induce expression of the *marR* operon (11). When MarR is incubated with increasing concentrations of NAFCs (naphthenic acid extracts from OSPW) (1.5-2 mg/ml), DNA binding is prevented, suggesting that compounds within this complex NA mixture is bound to MarR (Fig 7D). In general, there was a good agreement between the inducers of *marR* gene expression (11), or *in vivo* detection of NA, and the in vitro binding of NA in the EMSA biochemical assay.

### Expression of *marR-lux* and *atuA*-lux biosensors in *E. coli*

A further experiment to confirm the NA sensing and binding was to introduce the NA biosensor constructs into *E. coli*, and test if these strains can function as biosensors to detect NA exposure. Of the panel of *atuA* and *marR*-based biosensor constructs in this study, we predicted that some of these constructs would not be functional in *E. coli*, due to the lack of expression of the regulators from the low or medium strength constitutive *Pseudomonas* promoters (Fig 1, 3). This was confirmed as neither the atuR-L and marR-L third generation constructs in *E. coli* were functional to detect naphthenic acids mixture or individual compounds (data not shown). Since the high constitutive *Pseudomonas* promoter used to the control either the AtuR or MarR repressor was highly expressed in *E. coli* (Fig 3), we also introduced the atuA-H and marR-H third generation constructs into *E. coli* and tested their ability to detect NA.

The *E. coli* marR-H strain resulted in strong induction of the *marR* promoter during exposure to more complex NA compounds such as 5,6,7,8-Tetrahydro-2-naphthoic acid (THNA) and adamantane carboxylic acid (ACA),but also detected conventional NA compounds such as cyclohexane acetic acid (CHAA) and a mix of 9xNA compounds (Fig 8A). However, this strain was unable to detect the other compounds that could be detected by *Pseudomonas* (11), including cyclohexyl succinic acid (CHSA), decanoic acid and citronellic acid (Fig 8A). However, the *E. coli* biosensor responded with higher fold changes of gene expression than the *Pseudomonas* biosensor strains (11), when exposed to THNA (19-fold) or the diamondoid structure ACA (12-fold) (Fig 8A), suggesting that *E. coli* is more sensitive for detection of these compounds.

**Figure 8.**
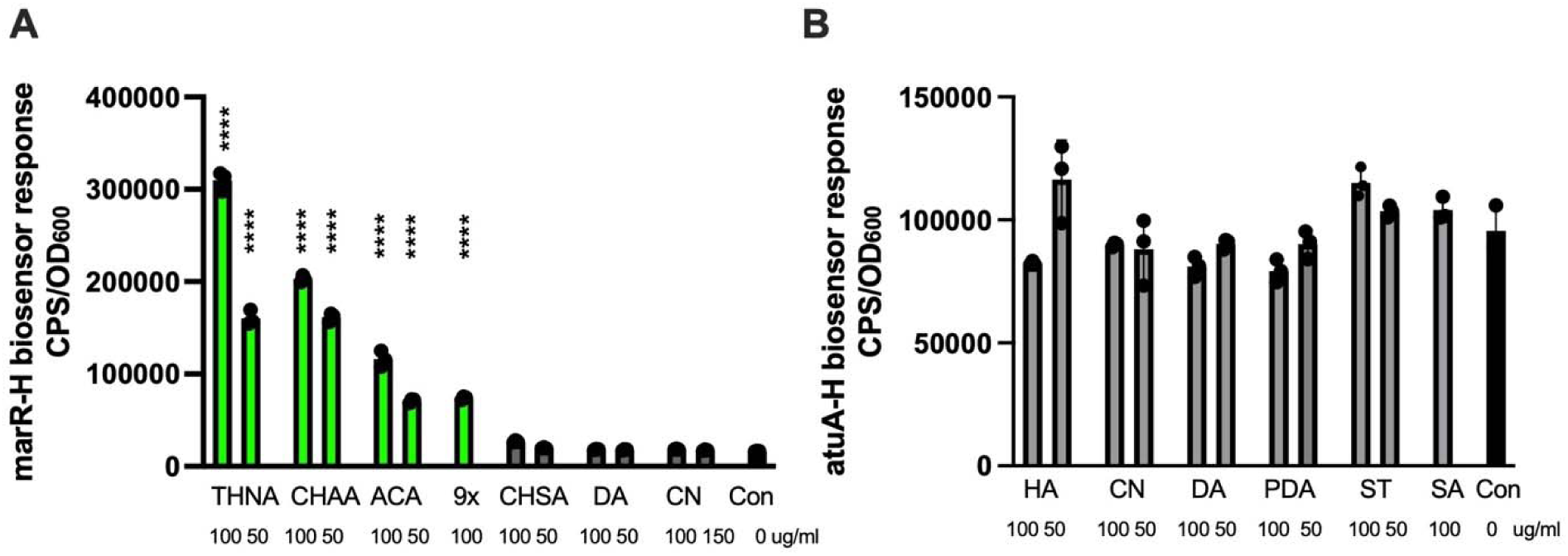
Expression of *marR* and *atuA*-based biosensors in *E. coli*. **A)** The third generation marR-H biosensor construct in *E. coli* was exposed to as 5,6,7,8-Tetrahydro-2-naphthoic acid (THNA), cyclohexane acetic acid (CHAA), adamantane carboxylic acid (ACA), 9x NA mix, decanoic acid (DA) and citronellic acid (CN) **B)** The third generation atuA-H biosensor construct in *E. coli* was exposed to hexanoic acid (HA), citronellic acid (CN), decanoic acid (DA), pentadecanoic acid (PDA), stearic acid (ST) or the commercial NA mixture (SA). All cultures were grown in diluted LB media (1/32). The gene expression responses were reported at the time points of maximal induction as CPS (counts per second) that was normalized by the OD_600_ (growth) with standard deviation values. All experiments were performed 4 times. A one-way ANOVA revealed a statistically significant difference between treatment groups (F(13,28) = 1158, p < 0.0001). A post hoc Dunnett’s test for multiple comparisons was conducted between each compound or mix to test significant relationships between each group (α = 0.05, n = 3). Treatment groups with significant results compared to the control are indicated with asterisks and green bars (**** p<0.0001). All gene expression values shown represent the average of triplicates with standard deviation and each experiment was performed 3 times.

In contrast, the *E. coli* atuA-H biosensor was not able to transcriptionally respond to multiple, fatty acid compounds that were detected by a *Pseudomonas* strain expressing this biosensor construct (Fig 8B). One possible reason may be related to the high copy number of pMS402 plasmid in *E. coli* and a low copy number in *Pseudomonas*, which will affect the overall levels of both the repressor protein and its’ target promoter. The copy number effect may not have allowed for sufficient AtuR to repress transcription of the *atuA* promoter in non-inducing conditions.

## Discussion

TetR and MarR-type repressor proteins are known to bind and respond to diverse small molecules, including fatty acids, and to regulate fatty acid degradation genes. This work has identified naphthenic acid and fatty acid compounds detected by MarR and AtuR, respectively, *in Pseudomonas* sp. OST1909. In the biosensor context, the MarR and AtuR repressors are involved in sensing and responding to compounds present in NAFCs and oil sands tailings water by inducing the expression of target genes (10, 23, 24). By varying the expression of the repressor proteins in a series of transcriptional *lux* reporter constructs, we were able to show decreased target gene expression when there was an increased level of repressor expression. The target promoters were shown to contain inverted repeat binding sites positioned within the promoter so that gene transcription via RNA polymerase would be blocked by repressor binding. Using purified, His-tagged AtuR and MarR proteins in EMSA assays, we demonstrated that the repressors bound their upstream DNA regions containing the operator sites.

The EMSA assay was also used as an approach to detect naphthenic acid binding to the repressors, which prevented DNA binding and inhibited the production of high molecular weight DNA-repressor protein complexes. Studies with the TfmR repressor, a TetR homolog, have also used the EMSA for the dual purpose of demonstrating promoter DNA binding but also binding to specific long chain fatty acid ligands and releasing from the promoter DNA (28). In our EMSA study, AtuR bound to a medium length fatty acid, decanoic acid, but not hexanoic acid, and also bound weakly to a commercial mixture of alkylated cyclopentane carboxylic acids and a large proportion of acyclic fatty acids (C5-C25, Z=0) (15, 16) that were isolated from a poorly defined hydrocarbon source. The AtuR-His_6_ tagged protein also bound to conventional NA compounds, such as CHPA, CPCA, ACA, THNA, which was unexpected because these latter compounds do not induce the expression of the *atu*A promoter (11). The *atu* operon appears to be an auxiliary fatty acid degradation gene cluster, distinct from the fatty acid degradation (*fad*) genes in the beta-oxidation pathway and is induced by compounds metabolized by these degradation genes (16–18).

The gene expression assay likely represents *in vivo* detection of fatty acids (saturated and branched) by AtuR, but the protein has a broader specificity of NA detection *in vitro* during a biochemical assay with the purified repressor protein and high concentrations of naphthenic acids. The reasons for this unexpected result are unclear, but the amount of NA required for *in vitro* binding was much higher than need to induce expression of the biosensor construct, and is not likely relevant to actual concentrations in the cell upon exposure to NA. In general, there was better agreement with the inducers of *marR* expression, and the NA compounds that were bound by MarR using the EMSA assay, such as fusaric acid, THNA, ACA, and CHAA. Further evidence that MarR is a naphthenic acid sensing protein was indicated by the expression of MarR in *E. coli*, and the ability of NA to induce expression of the *marR-lux* transcriptional fusion from the marR-H biosensor constructs.

Collectively, these data provide evidence for AtuR and MarR as repressors and fatty acid or naphthenic acid sensing proteins, which also provides a better mechanistic understanding of the gene expression patterns and their potential application as a biosensor technology. The naphthenic acids biosensors are a novel monitoring technology that can be used to detect NA in oil sands generated wastewater that is stored on site. By comparing different bacterial hosts for the NA biosensor constructs, we demonstrated that *E. coli* is more sensitive for the detection of ACA, a recalcitrant, diamondoid structure and therefore may have future applications in ACA monitoring or bioremediation studies.

## Acknowledgments

This work was supported by an NSERC Discovery Grant and by Genome Canada through a Large Scale Applies Research Project (LSARP) grant, project number 18207, in partnership with Genome Alberta and Genome Quebec. Co-funding was provided by the Government of Alberta through an Alberta Innovates grant and Jobs, Economy and Innovation funding.

